# Non-monotonic effects of GABAergic synaptic inputs on neuronal firing

**DOI:** 10.1101/2021.12.07.471426

**Authors:** Aghil Abed Zadeh, Brandon D. Turner, Nicole Calakos, Nicolas Brunel

## Abstract

GABA is generally known as the principal inhibitory neurotransmitter in the nervous system, usually acting by hyperpolarizing membrane potential. However, GABAergic currents sometimes exhibit non-inhibitory effects, depending on the brain region, developmental stage or pathological condition. Here, we investigate the diverse effects of GABA on the firing rate of several single neuron models, using both analytical calculations and numerical simulations. We find that GABAergic synaptic conductance and output firing rate exhibit three qualitatively different regimes as a function of GABA reversal potential, *E*_GABA_: monotonically decreasing for sufficiently low *E*_GABA_ (inhibitory), monotonically increasing for *E*_GABA_ above firing threshold (excitatory); and a non-monotonic region for intermediate values of *E*_GABA_. In the non-monotonic regime, small GABA conductances have an excitatory effect while large GABA conductances show an inhibitory effect. We provide a phase diagram of different GABAergic effects as a function of GABA reversal potential and glutamate conductance. We find that noisy inputs increase the range of *E*_GABA_ for which the non-monotonic effect can be observed. We also construct a micro-circuit model of striatum to explain observed effects of GABAergic fast spiking interneurons on spiny projection neurons, including non-monotonicity, as well as the heterogeneity of the effects. Our work provides a mechanistic explanation of paradoxical effects of GABAergic synaptic inputs, with implications for understanding the effects of GABA in neural computation and development.

**Author summary:** Neurons in nervous systems mainly communicate at chemical synapses by releasing neurotransmitters from the presynaptic side that bind to receptors on the post-synaptic side, triggering ion flow through ion channels on the cell membrane and changes in the membrane potential of the post-synaptic neuron. Gamma-aminobutyric acid (GABA) is the principal neurotransmitter expressed by inhibitory neurons. Its binding to GABAergic ionotropic receptors mainly causes a flow of chloride ions across the membrane, and typically hyperpolarizes the post-synaptic neuron, resulting in firing suppression. While GABA is canonically viewed as an inhibitory neurotransmitter, non-inhibitory effects have been observed in early stages of development, in stress-related disorders, and in specific parts of brain structures such as cortex, cerebellum and hippocampus [1–4]. Here, we employ analytical and computational approaches on spiking neuronal models to investigate the mechanisms of diverse effects of GABAergic synaptic inputs. We find that in addition to monotonically excitatory or monotonically inhibitory effects, GABAergic inputs show non-monotonic effects, for which the effect depends on the strength of the input. This effect is stronger in the presence of noise, and is observed in different models both at the single cell, and at the network level. Our findings provide a mechanistic explanation of several paradoxical experimental observations, with potential implications for neural network dynamics and computation.

## Introduction

GABA is the principal inhibitory neurotransmitter in the nervous system. In adult animals, GABA usually suppresses action potentials in target neurons by hyperpolarizing the membrane potential. This hyperpolarization is mediated by GABA receptor channels, that are permeable to Cl^−^ and 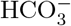 ions. The flux of these ions usually causes a decrease in membrane potential upon GABA channels opening [5, 6].

GABAergic synapses are canonically viewed as inhibitory. However, multiple studies have found that GABA can have non-inhibitory effects. For instance, excitation mediated by GABA plays an important role in early phases of development and neural integration during neurogenesis. It has been shown that this excitatory effect is caused by a depolarizing effect of GABA on neurons [1, 7–10]. Other studies suggest that pathological conditions of stress or trauma can lead to excitatory effects of GABA [2, 11]. Moreover, even in healthy adult animals, GABA mediates excitatory effects in certain brain regions. This excitatory effect of GABA has been observed in several brain regions including cortex, basal ganglia, thalamus and cerebellum [4, 12–16]. In addition to excitatory effects of GABA, other studies have shown non-monotonic effects of GABA, suggesting that GABAergic synaptic currents produce excitation or inhibition based on their strength. In hippocampal interneurons, it has been shown that changing tonic GABA conductance by varying extra-cellular GABA concentration affects neuronal firing in a non-monotonic way [17]. In the striatum, one study shows that bidirectional optogenetic manipulations (inhibition and excitation) of fast spiking interneurons (FSIs) cause spiny projection neurons (SPNs) population activity inhibition [18, Fig 1], suggesting that SPNs activity may depend on FSI GABAergic inputs in a non-monotonic way. In the non-monotonic cases, small GABAergic inputs promote neural firing while large currents have an inhibitory effect. Non-inhibitory GABAergic inputs significantly influence network dynamics too. Experimental studies and computational models show effects of such depolarizing GABA currents on neural synchrony and rhythmicity [19–21]. Understanding the mechanisms of such effects in neural dynamics is central to unravel their role on neural computation and plasticity and combat related diseases.

**Fig 1.**
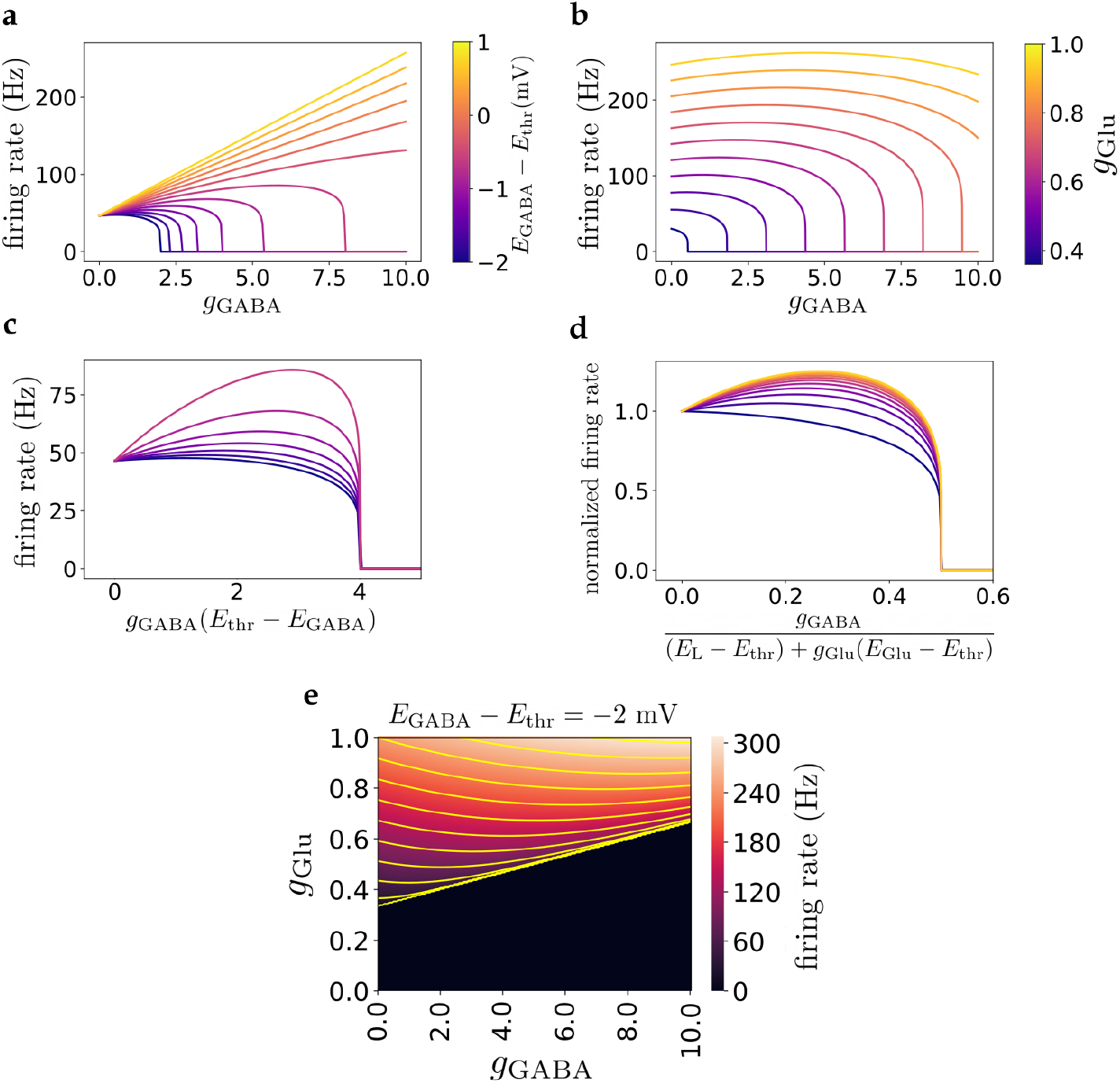
Effect of *g*_GABA_ on firing rate in the deterministic LIF model. **a)** Firing rate *ν*, as a function of GABAergic conductance for different GABA reversal potentials (*g*_Glu_ = 0.4). When GABA reversal potential is below and close to firing threshold, *E*_thr_, the firing rate has a non-monotonic dependence on *g*_GABA_. **b)** Firing rate vs *g*_GABA_ for different *g*_Glu_, (*E*_GABA_ − *E*_thr_ = −3 mV). The non-monotonicity appears when the glutamatergic conductance is sufficiently large. **c)** Firing rate vs GABA conductance rescaled with *E*_thr_ − *E*_GABA_, for *E*_thr_ > *E*_GABA_. For GABA reversal potentials smaller than the threshold, complete inhibition occurs for large GABA conductances. The onset of zero-firing scales with 1/(*E*_thr_ − *E*_GABA_) as shown by the collapsed curves (same color code as above). **d)** Firing rate vs rescaled *g*_GABA_. The collapsed curves show how the zero-firing onset scales with *g*_Glu_ (same color code as above). **e)** Heatmap of firing rate as a function of GABAergic and glutamatergic conductances for *E*_GABA_ − *E*_thr_ = −2 mV. The yellow curves are contour lines of constant firing rates.

In this paper, we investigate potential mechanisms of different GABA effects in neuronal and circuit models. We show how changing GABA reversal potential affects the firing rate of neurons. In particular, we show that GABA can be inhibitory, non-monotonic or excitatory depending on neuron’s reversal potential for GABA. The non-monotonic regime is observed when GABA reversal potential is below, but close to the neuron’s firing threshold. Using analytical calculations in a leaky integrate and fire (LIF) conductance-based model, we provide a phase diagram of the dynamics and analyze it in the presence of input noise, showing that noisy input expands the non-monotonic regime. We also investigate a more realistic model that describes more accurately the electrophysiological properties of SPNs in the striatum, to check the robustness of our model. Finally, we study a network model of striatum to explain several observed paradoxical effects of GABAergic currents from striatal FSIs. The motivation to focus on striatum is the observation of multiple effects of FSIs on their SPN targets that cannot be explained by GABA actions that are solely inhibitory or solely excitatory [14, 18, 22]. Furthermore, it is known that striatal FSIs influence behavioral outputs, and their dysfunction is implicated in neurological disease and movement disorders such as dystonia and Tourette syndrome [23–27]. We use network simulations and computational analysis to provide possible insights into non-monotonic and heterogeneous effects of GABAergic inputs on their targets. These analyses can be used to reconcile several experimental findings and to provide a framework for understanding the role of GABA in network dynamics and plasticity in different brain regions and developmental stages.

## Materials and methods

We use three different models to study the effects of GABAergic synaptic inputs on neuronal firing rate. We start by analyzing a single-cell conductance-based LIF model which is analytically tractable. We then expand our computational analysis on a more realistic neuronal model. Finally, we implement and analyze a micro-circuit model of striatum.

### Leaky integrate-and-fire model

This model describes the dynamics of the membrane potential which is given by a current balance equation, with capacitive and leak currents, together with conductance-based GABAergic and glutamatergic synaptic currents. Additionally a stochastic Gaussian noise is used to study the effect of noisy inputs. The model can be described by a small number of variables, and at the same time captures different effects of GABAergic conductance on firing activity. The evolution of the membrane potential follows

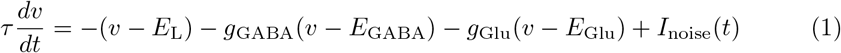

when the membrane potential is smaller than a threshold, *v* < *E*_thr_. Spikes are emitted whenever *v* = *E_thr_*, after which the voltage is reset instantaneously to *E*_reset_. We use standard model parameters, set to be in the range of experimentally observed values and shown in Table 1, except when specified otherwise. However, the space of glutamatergic and GABAergic conductances is explored systematically. The reported results are qualitatively robust to changes of single neuron variables in a wide range of such parameters. Note that in Eq (1), *g*_GABA_ and *g*_Glu_ are dimensionless, as both sides of the equation have been divided by the leak conductance *g*_L_. The last term in the right hand side of Eq (1) is a noise term, 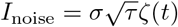 in which *ζ* is a Gaussian white noise with unit variance. We first analyze the dynamics in the absence of noise (*σ* = 0) and then investigate the effects of noise on the dynamics. We study two different classes of noise. First, we consider additive noise in which *σ* is a constant, independent of membrane potential and conductances. Second, we study noise that represents fluctuations caused by Poisson firing of presynaptic neurons, in which case, *σ* depends on several model parameters and membrane potential.

### EIF-Kir model

While most of our analysis is based on the model of Eq (1), we also analyze a more realistic neuronal model to show the qualitative robustness of our results. In this model, as represented in Eq (2), two more currents are added: an exponential current (*I*_Exp_) [28] that represents in a simplified fashion the fast sodium currents near the spiking threshold, and an Inward-Rectifier potassium current (*I*_Kir_) that captures the nonlinear dependence of effective conductance of membrane potential. The dynamics of this EIF-Kir model is given by

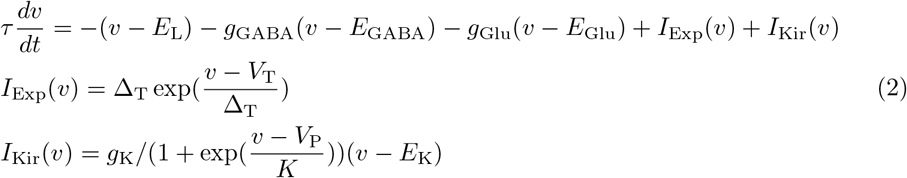

The parameters of the model are chosen such that it reproduces sub-threshold (current-voltage relation) and supra-threshold (current-firing rate relation) properties of a typical spiny projection neuron in striatum (see Fig 5c-d and refs [29]). Model parameters are shown in Tables 1 & 2 unless specified otherwise. As in the LIF model, all conductances, including *g*_K_, are normalized by the leak conductance and thus dimensionless. In this model, there is no hard spike threshold, rather the membrane potential resets when the voltage diverges to infinity due to the exponential spike-generating current. To calculate V-I and f-I curves, we add a constant current of form *I*_const_ = *I*/*g*_L_ to Eq (1b) in which *I* is in the units of pA and we set *g*_L_ = 5nS.

**Table 1.** LIF model parameters

| parameter | description | value |
| --- | --- | --- |
| $\tau$ | membrane time constant | 20 ms |
| $E_L$ | leak reversal potential | -80 mV |
| $g_{\text{GABA}}$ | GABA conductance (normalized by $g_L$) | variable |
| $g_{\text{Glu}}$ | glutamate conductance (normalized by $g_L$) | variable |
| $E_{\text{GABA}}$ | GABA reversal potential | variable |
| $E_{\text{Glu}}$ | glutamate reversal potential | 0 mV |
| $E_{\text{thr}}$ | spike threshold | -60 mV |
| $E_{\text{reset}}$ | reset potential | -70 mV |
| $I_{\text{noise}}$ | noise term | $\sigma\sqrt{\tau}\zeta(t)$ |

**Table 2.**
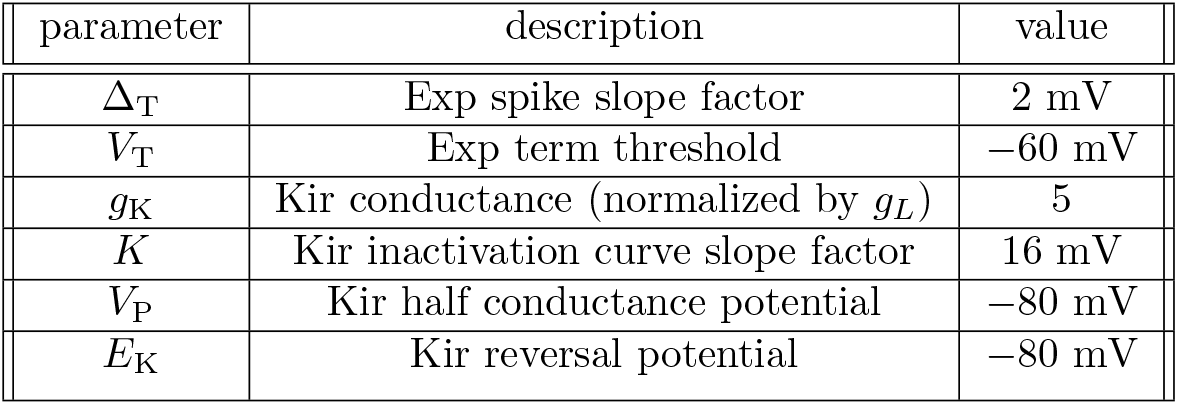
EIF-Kir model parameters

| parameter | description | value |
| --- | --- | --- |
| $\Delta_T$ | Exp spike slope factor | 2 mV |
| $V_T$ | Exp term threshold | -60 mV |
| $g_K$ | Kir conductance (normalized by $g_L$) | 5 |
| $K$ | Kir inactivation curve slope factor | 16 mV |
| $V_P$ | Kir half conductance potential | -80 mV |
| $E_K$ | Kir reversal potential | -80 mV |

### Population-level model

We used a micro-circuit model of a striatal network to demonstrate paradoxical effects of GABA at the network level. Single neurons were described by Eq (1) with only leak, GABAergic and glutamatergic currents. There are three types of neurons in the model: direct and indirect spiny projection neurons (dSPN & iSPN) and fast spiking interneurons (FSI). The network is composed of *N* = 1000 neurons with 2% FSI, 49% dSPN and 49% iSPN, approximating a simplified micro-circuit of ~ 0.01 mm^3^ of mouse striatum [29, 30], but not accounting for other interneuron cell types as FSIs provide the major GABAergic inputs to SPNs [29, 31]. FSIs have a GABA reversal potential of −80 mV and provide feedforward GABAergic currents to SPNs. The glutamatergic conductances are taken for simplicity to be driven by independent Poisson inputs convoluted with an exponential kernel and a frequency of 1000 Hz, approximating the total cortical input to a striatal neuron. The exponential kernel has a decay time constant of 5.6 ms (similar to [32]). The mean value of this Poisson excitatory input is chosen to reproduce the desired firing rates in the model. Neurons are connected by GABAergic synapses in which a spike in presynaptic neuron produces a post-synaptic conductance (PSC) of form

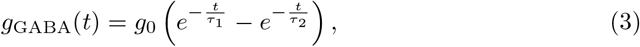

with a decay time of *τ*_1_ = 20 ms and a rise time of *τ*_2_ = 1.5 ms [33], shown in Fig 6b. The connectivity probabilities and strengths are inferred from experimental studies [31, 33]. The connection probabilities *P_ij_* and strengths *G_ij_* of synaptic connections from population *j* to population *i* (from column *j* to row *i*), where indices 1, 2, 3 stand for FSIs, dSPNs and iSPNs respectively, are

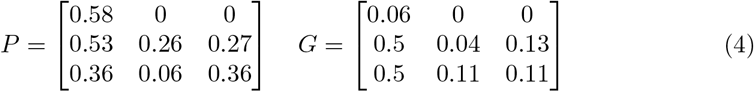

Note that *G* entries are normalized and dimensionless, being the ratio of synaptic to leak conductances, similar to *g*_GABA_ in Eq (2). The average total normalized conductance from population *j* with *N_j_* neurons, firing at a mean rate of *ν_j_*, to population *i* is *G_ij_P_ij_N_j_v_j_* (*τ*_1_ − *τ*_2_). In simulations, the first 100 ms are removed from any analysis to avoid transient effects.

## Results

### Effects of GABA on the deterministic LIF model

To gain insight into the mechanisms of different effects of GABA, we start by analyzing the simplest possible model, i.e. the one-dimensional LIF model described Eq (1), with no added noise and constant synaptic conductances. The simplicity of this model allows us to rigorously characterize the different regimes of GABA effects on neuronal firing rate through analytical calculations. The dynamics of the membrane potential is in this case deterministic and can be rewritten as

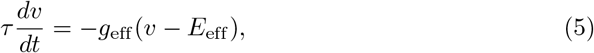

where the effective input conductance *g*_eff_ (relative to the leak) and the effective reversal potential *E*_eff_ are given by

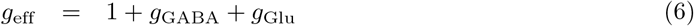

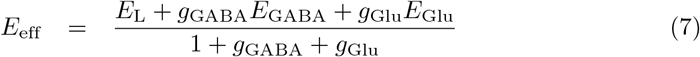

According to Eq (5), the membrane potential relaxes exponentially to an effective potential, *E*_eff_, with an effective time constant *τ*_eff_ = *τ*/*g*_eff_. If *E*_eff_ is greater than *E*_thr_, the neuron spikes with a non-zero firing rate *v*, given by

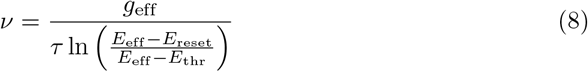

Fig 1 shows examples of how the firing rate depends on *g*_GABA_ for different values of *g*_Glu_ and *E*_GABA_. As shown in Fig 1a,b, the firing rate vs GABA conductance curves can be decreasing, non-monotonic or increasing, depending on *E*_GABA_ and *g*_Glu_. When *E*_GABA_ > *E*_thr_, GABAergic current is always excitatory. When *E*_GABA_ < *E*_thr_, large *g*_GABA_ leads to inhibition; however, it can be excitatory for smaller values of *g*_GABA_.

When *E*_GABA_ < *E*_thr_, large enough *g*_GABA_ silences the neuron. The GABAergic conductance for which the firing stops, 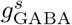, can be calculated using the condition *E*_eff_ = *E*_thr_. This leads to

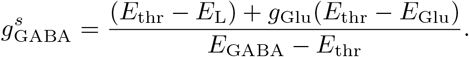

Fig 1c,d shows the collapse of zero-firing onset when *g*_GABA_ is scaled appropriately with respect to *E*_GABA_ and *g*_Glu_. This onset increases with *g*_Glu_ and *E*_GABA_ and diverges to infinity as *E*_GABA_ → *E*_thr_.

The occurrence of the different regimes of GABA effects on firing rate can be understood by analyzing Eq (8). In this equation, *g*_GABA_ acts on the firing rate in two ways: Through its effects on *g*_eff_, and on *E*_eff_. Increasing GABA conductance increases *g*_eff_, which tends to increase the firing rate due to the decrease in effective membrane time constant of the neuron, while the effect of GABA on *E*_eff_ depends on the GABA reversal potential. The competition between these two effects can be analyzed by computing the derivative of the firing rate with respect to the GABA conductance,

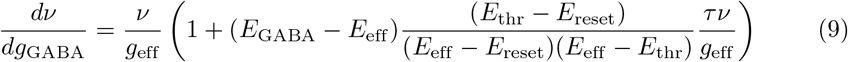

where the first and second terms within the parentheses represent the effects of *g*_eff_ and Eeff on the firing rate, respectively. The sign of this derivative determines whether GABAergic conductance is inhibitory or excitatory. The sufficient conditions for non-monotonic effect can be summarized as:

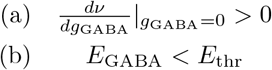

Condition (a) makes GABA excitatory for small *g*_GABA_ and the condition (b) makes GABA inhibitory for large *g*_GABA_. These conditions can be met when the term *E*_eff_ − *E*_GABA_ in Eq (9) is positive but not too large, which means *E*_eff_ should be above and close to *E*_GABA_. The critical value of *E*_GABA_ that separates the inhibitory and non-monotonic regime, 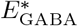, can be obtained from Eq (9) setting *dν*/*dg*_GABA_ = 0 at *g*_GABA_ = 0:

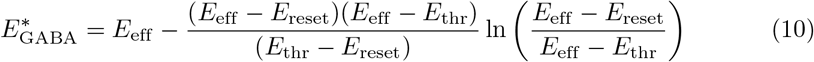

For values of *g*_Glu_ that are just above the threshold for firing (given by *g*_Glu_ = (*E*_thr_ − *E*_L_)/(*E*_Glu_ − *E*_thr_) ≈ 0.33 here), *E*_eff_ ≈ *E*_thr_. This leads to 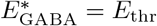. On the other hand, for large *g*_Glu_, *E*_eff_ ≫ *E*_thr_, which leads to 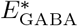 converging to (*E*_reset_ + *E*_thr_)/2. This result shows that in the deterministic LIF model, the non-monotonic effect can be observed when *E*_GABA_ is between (*E*_reset_ + *E*_thr_)/2 and *E*_thr_. Using model parameters of Table 1, non-monotonic behavior can be present whenever −5 mV< (*E*_GABA_ − *E*_thr_) < 0.

These results are shown in the phase diagram of Fig 2a. This phase diagram separates the *E*_GABA_ − *g*_Glu_ plane into five regions: For sufficiently high *g*_Glu_ (above the horizontal line in Fig 2a), GABA is always inhibitory for 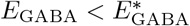 (Fig 2b); The GABA effect is non-monotonic for 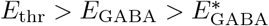 (Fig 2c); GABA is always excitatory for *E*_thr_ < *E*_GABA_ (Fig 2d). Finally, for low *g*_Glu_ (below the horizontal line in Fig 2a), the neuron either remains silent for all GABA conductances for *E*_GABA_ < *E*_thr_, or starts firing with an increasing rate beyond a critical value of the GABA conductance for *E*_GABA_ > *E*_thr_. Fig 2a also shows how the slope of the firing rate vs *g*_GABA_ curve at zero GABA conductance, i.e. 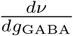, depends on both glutamatergic conductance and GABA reversal potential.

**Fig 2.**
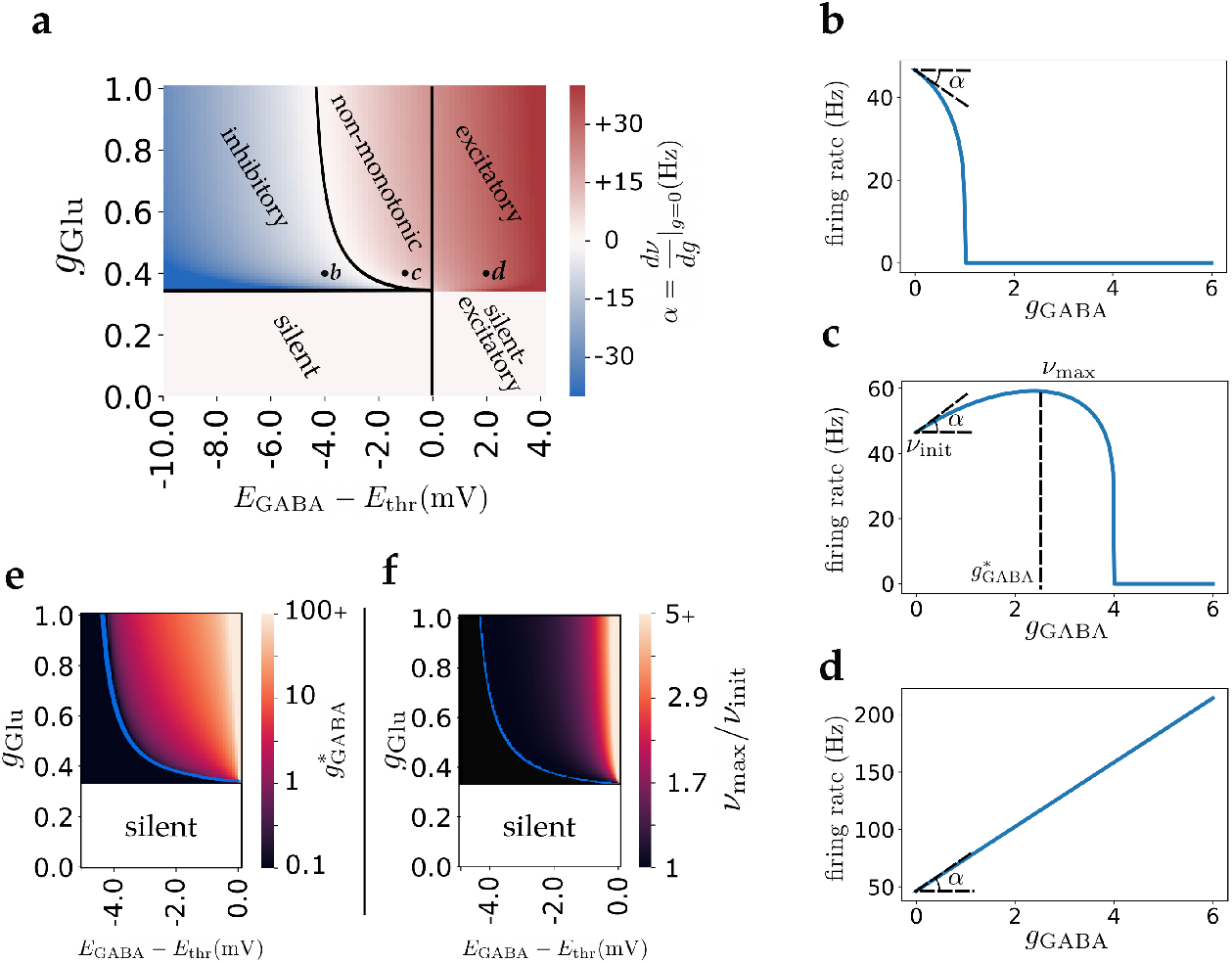
GABA effect phase diagram in the deterministic LIF model. **a)** Phase diagram of GABA effects. The colorbar indicates the initial slope of firing curves with respect to *g* ≡ *g*_GABA_ as shown in (b-d). b, c, d marked in the phase diagram correspond to three example firing rate curves below. **b,c,d)** As *E*_GABA_ increases, GABA effect changes from inhibitory to excitatory with a non-monotonic region in between. In this region, GABA conductance below a certain threshold, 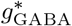, has an excitatory effect while large GABA conductances have an inhibitory effect. **e)** 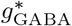 (GABA conductance that maximizes output firing rate, shown in part c). **f)** Ratio of maximum firing rate to firing rate at *g*_GABA_ = 0 (*ν*_max_/*ν*_init_ shown in part c), as a measure of non-monotonic effect strength. In **e** & **f**, the non-monotonic regime is on the right of the blue curves and colorbars are in logarithmic scale.

Fig 2e,f provide additional characterizations of the non-monotonic regime. Fig 2e shows how the GABA conductance that maximizes the firing rate depends on *E*_GABA_ and *g*_Glu_. The colorbar represents 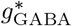 as shown in Fig 2c example. Fig 2f shows the strength of the non-monotonic effect, i.e. the ratio between maximal firing rate *ν*_max_ and firing rate in the absence of GABA *ν*_init_ in the non-monotonic region. Here *ν*_init_ = *ν*(*g*_GABA_ = 0) and *ν*_max_ = max(*ν*(*g*_GABA_)). This ratio converges to 1 when 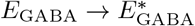, and diverges in the limit that *E*_GABA_ → *E*_thr_.

To conclude, in the deterministic LIF model, there is a region of parameters for which GABA has a non-monotonic effect on neuronal firing rate, when *E*_GABA_ is below but sufficiently close to *E*_thr_. However, this effect is weak except in a narrow range of *E*_GABA_ close to threshold, as shown in Fig 2f.

### Input noise expands the non-monotonic regime

So far, we have investigated the effect of constant deterministic GABAergic synaptic inputs. In real neurons, the synaptic currents are noisy for multiple reasons, including stochastic vesicle release and channel opening. In addition, presynaptic neuronal firing can often exhibit a large degree of irregularity, which is typically approximated by a Poisson process. The total synaptic currents to a neuron can then be described by the sum of their temporal mean, and stochastic fluctuations around the mean. Here, we consider two simplified models for noise. First, we use a noise of a constant amplitude, that is independent of the glutamatergic and GABAergic conductances. This simplification facilitates calculations and provides insights into the effect of noise. Second, we consider noise as originating from fluctuations in glutamatergic and GABAergic conductances. This is a more realistic assumption and is discussed in the latter part of this section.

We start by investigating the effects of white noise with a fixed amplitude on how GABA affects neuronal firing rate. The firing rate of the neuronal model of Eq (1) in the presence of noise is given by [34, 35]:

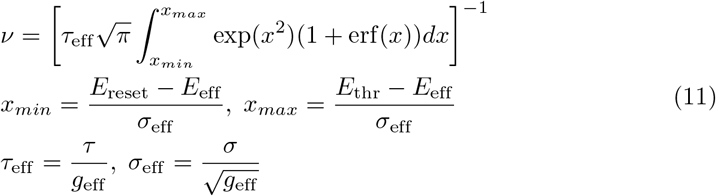

We start by considering the case when *σ* is independent of other parameters. Fig 3a shows the firing rate as a function of *g*_Glu_ for different values of *σ* (color-coded) and *g*_GABA_ = 0. When there is no noise, *σ* = 0, the firing rate is zero below a certain threshold of *g*_Glu_, increases sharply near this threshold and finally converges to a linear relation (note that the lack of saturation is due to our choice of setting the refractory period to zero). As *σ* increases, the curves become smoother with the neuron firing for sub-threshold values of *g*_Glu_, due to fluctuations of membrane potential. This effect allows the neuron to have low firing rates for a wider range of *g*_Glu_, compatible with the observed low firing rate *in vivo* in many brain regions (~ 1 Hz). The effect of noise on the different regimes of how GABA affects neuronal firing rate are illustrated in Fig 3b,c. Both of these figures show firing rate as a function of GABAergic conductance for different values of noise. In Fig 3b all curves receive the same *g*_Glu_, so the noise increases the firing rate and non-monotonic effect simultaneously. In Fig 3c, *g*_Glu_ is chosen such that initial firing rate is the same (*ν*_init_ = 1 Hz) for different noise levels. This figure also shows that increasing *σ* makes the curves peak value larger, thus the non-monotonic effect becomes stronger with increasing noise level.

**Fig 3.**
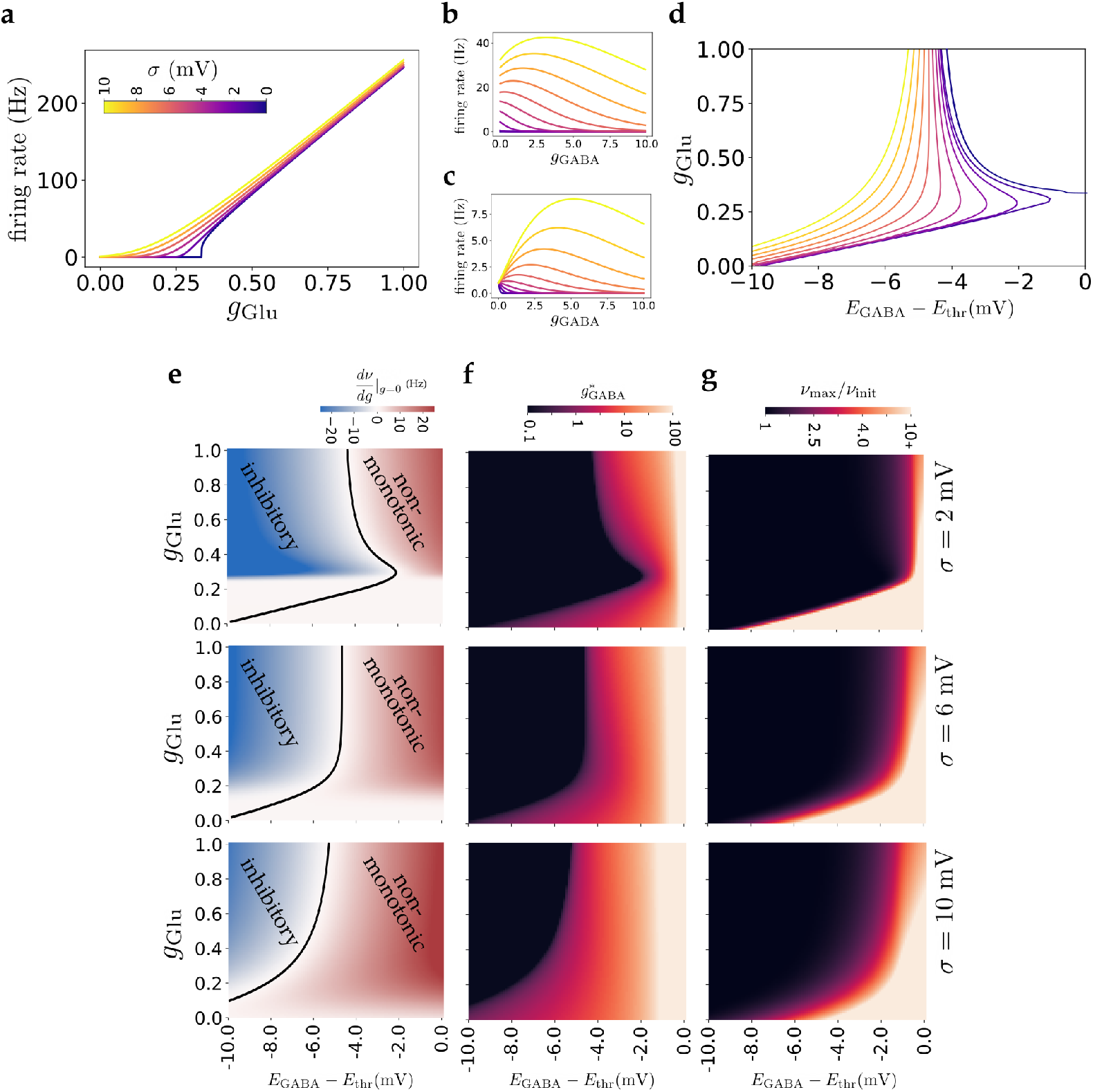
Effect of current-based input noise. **a)** Transfer functions for different noise levels. Increasing noise smooths the transfer function for low firing rates and results in firing for subthreshold values of *g*_Glu_. **b)** Effect of *g*_GABA_ on firing rate for different noise levels (*g*_Glu_ = 0.25, *E*_GABA_ − *E*_thr_ = −5 mV). Increasing noise level, while keeping *g*_Glu_ constant, increases both the firing rate and the non-monotonic effect. **c)** Effect of *g*_GABA_ on firing rate for different noise levels (*v*_init_ = 1 Hz, *E*_GABA_ − *E*_thr_ = −5 mV). Here *g*_Glu_ is modified for different values of *σ* to get same initial firing rate. Increasing noise in this case also strengthens the non-monotonic effect. **d)** Boundary between the non-monotonic and inhibitory regions of phase diagrams for different noise levels. Increasing noise extends the non-monotonic region for all values of *g*_Glu_, with a more drastic change for smaller *g*_Glu_. **e)** Examples of phase diagrams similar to that of Fig 2 for different noise levels. Introducing noise removes the silent region and enlarges the non-monotonic region. **f)** Similar phase diagrams as in part e, showing 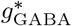. **g)** Similar phase diagrams as in part e, showing the ratio of max firing rate to initial firing rate.

For non-zero noise, the neuron exhibits a non-zero firing rate for any value of *g*_Glu_. Thus, the silent region of the phase diagram in Fig 2 disappears. The boundary between non-monotonic and excitatory regions remains unchanged at *E*_GABA_ = *E*_thr_. In the case that *E*_GABA_ > *E*_thr_, for large *g*_GABA_, *E*_eff_ → *E*_GABA_ > *E*_thr_ and *g*_eff_ → inf; using Eq (8), the firing rate diverges and GABAergic currents are always excitatory. However, noise changes the boundary that separates the inhibitory and non-monotonic regions. Fig 3d shows how the boundary between inhibitory and non-monotonic regions of the phase diagram changes as *σ* increases, while Fig 3e shows examples of the phase diagram for several values of *σ*. Increasing *σ* expands the non-monotonic region, with a stronger effect for smaller values of *g*_Glu_. This effect can be shown mathematically using Eq (11) in the limit of large *σ* and small *g*_Glu_ and *g*_GABA_. In this case, as the limits of integral tend to zero, the integrand can be approximated by 1, resulting in:

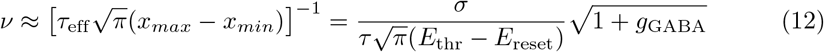

In this limit, the firing rate is an increasing function of *g*_GABA_ which shows an excitatory effect for small GABAergic conductances, as expected in the non-monotonic regime. As a result, in the presence of strong fluctuating noise, non-monotonic effect of GABA input is present for a wide range of *E*_GABA_. Fig 3f shows how the GABA conductance 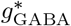 that maximizes firing rate depends on other parameters, while Fig 3g shows how the strength of the non-monotonic effect depends on *E*_GABA_ and *g*_Glu_. In particular, this figure shows that there is a much larger range of these parameters for which the non-monotonic effect is strong (i.e. ratio of maximal to initial firing rate significantly higher than 1).

So far, we have investigated the effect of noise whose amplitude *σ* is a constant and is independent of conductances. This approach reveals the qualitative effect of noise; however, here we extend our analysis by considering a more realistic representation of noise. The main source of noise *in vivo* is thought to have a synaptic origin, and be due to both irregular firing of presynaptic neurons and stochastic release of synaptic vesicles. This stochasticity causes fluctuations that depend on the strengths of inputs, i.e. stronger inputs cause higher fluctuations. Here, we assume instantaneous synaptic conductances that are a sum of delta functions:

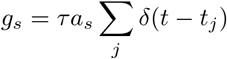

in which *s* ∈ {Glu, GABA}, presynaptic spikes are generated at times *t_j_* leading to an instantaneous change of magnitude *a_s_*(*v_s_* − *v*) in neuron’s membrane potential. A neuron receives synaptic inputs of type *s* ∈ {Glu, GABA}from *K_s_* other neurons. In the limit of *K_s_* ≫ 1, *a_s_* ≪ 1, assuming uncorrelated Poisson firing of presynaptic neurons, synaptic inputs can be approximated by (e.g. [35–37]):

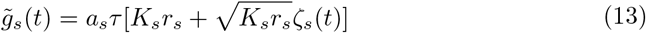

in which *s* can be Glu or GABA, *g_s_* = *a_s_τK_s_r_s_* is the average synaptic input, *r_s_* is the average firing rate of connected neurons and *ζ_s_* is a white Gaussian noise with zero mean and unit variance. Assuming independent glutamatergic and GABAergic synaptic inputs, the equation for the membrane voltage reduces to Eq (1) with noise term parameter:

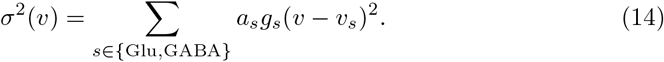

Here, as the noise term depends on the membrane potential v, it is a multiplicative noise. It can be shown, however, [36] that this term can be approximated by substituting *v* by *E*_eff_, *σ*(*v*) ≈ *σ*(*E*_eff_), which makes the noise term independent of *v* and easier to analyze. Using this approximation, we can obtain firing curves by applying Eq (11). Fig 4a shows examples of firing rate vs GABA conductance curves obtained using this approximation. In this case, we set the individual synaptic strengths *a*_Glu_ = *a*_GABA_ = *a*. A non-monotonic effect of GABA can be observed with this model, similar to the constant noise case. The strength of noise in this model can be modulated by the value of a, as can be seen in Eq (14). Fig 4b-d provide phase diagrams of GABA effects for three different values of a. As can be seen in Fig 4, increasing noise expands the non-monotonic regime. This expansion is more significant for higher values of *g*_Glu_. This is in contrast to the effect of constant noise, shown in Fig 3e, in which noise predominantly expands this regime for lower values of *g*_Glu_. This effect is due to the fact that in the conductance-dependent noise case, the noise term depends on the value of *g*_Glu_ and *σ* vanishes as conductances go to zero, such that the noise has little effect for low values of *g*_Glu_. Overall, the results of both current-based and conductance-based noise suggest that non-monotonic effects become stronger as input noise increases.

**Fig 4.**
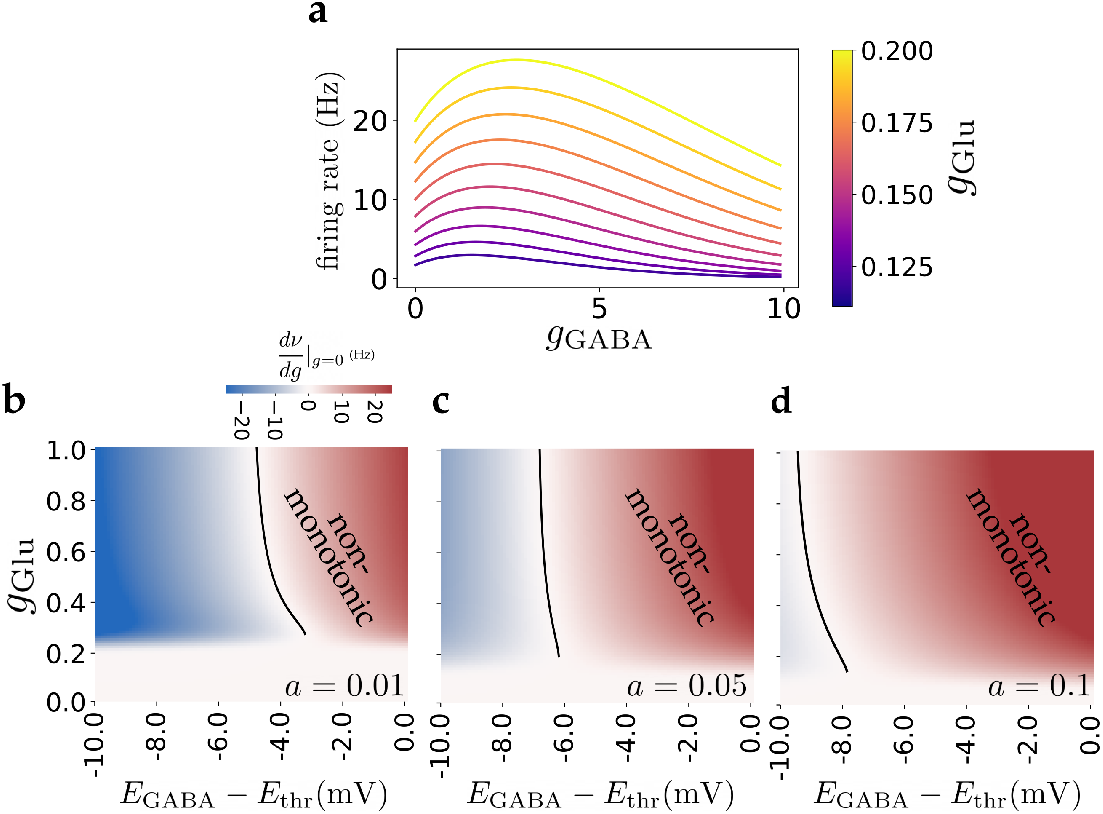
Effect of conductance-dependent input noise. **a)** Examples of firing rate vs GABAergic conductance curves, showing non-monotonic effect. Different colors represent different glutamatergic inputs. Here, *a* = 0.1 and *E*_GABA_ − *E*_thr_ = −5 mV. **b,c,d)** Phase diagrams of GABA effect, similar to Fig 3e, for noise parameters *a* = 0.01, 0.05, 0.1 respectively. Non-monotonic regime extends with increasing noise.

### Non-monotonic effects in a more realistic neuronal model

To check that GABA effects are not specific to the LIF model, we analyze the dynamics of a neuronal model described by Eq (2), in which two more currents are added: An exponential current (Exp) that influences the near threshold dynamics and spike generation [28], and an inward-rectifier potassium current (Kir) that accounts for non-linear dependence of membrane conductivity as a function of membrane potential (see e.g. [38] chapter 4.4.3). Fig 5b shows an example simulation of this model in response to a constant input conductance. The Exp term changes the near threshold dynamics and the Kir current reproduces the non-linear dependence of voltage on current, as observed in several neuronal types, including striatal neurons [39] (see Fig 5c). In this model, there is no hard spiking threshold as in Eq (1), rather the Exp term generates spikes whenever the voltage gets sufficiently close to V_T_, the Exp term threshold, so that the Exp term leads to divergence of the voltage. Fig 5e shows the effect of GABAergic currents for different GABA reversal potentials. As shown in the figure, the non-monotonic effect is observed when the GABA reversal potential is close to *V*_T_. For *E*_GABA_ far from *V*_T_, the GABA is effectively inhibitory or excitatory, qualitatively similar to that of Eq (1). The non-monotonic curves of Fig 5e are slightly different compared to those of Fig 1a mainly because of the Exp term. The effect of the exponential spike-generating current on the non-monotonic regime can be seen in Fig 5f. As Δ_*T*_ in Eq (2) increases (and therefore spike generation becomes less sharp), the influence of the exponential term grows. For low values of *E*_GABA_, the non-monotonic regime becomes inhibitory, while for higher values of *E*_GABA_, the excitatory regime becomes non-monotonic. Thus, increasing Δ_*T*_ effectively shifts the non-monotonic regime to higher values of GABA reversal potential. Analysis of the model with the Exp term but no Kir current produces very similar curves as in Fig 5e (not shown).

**Fig 5.**
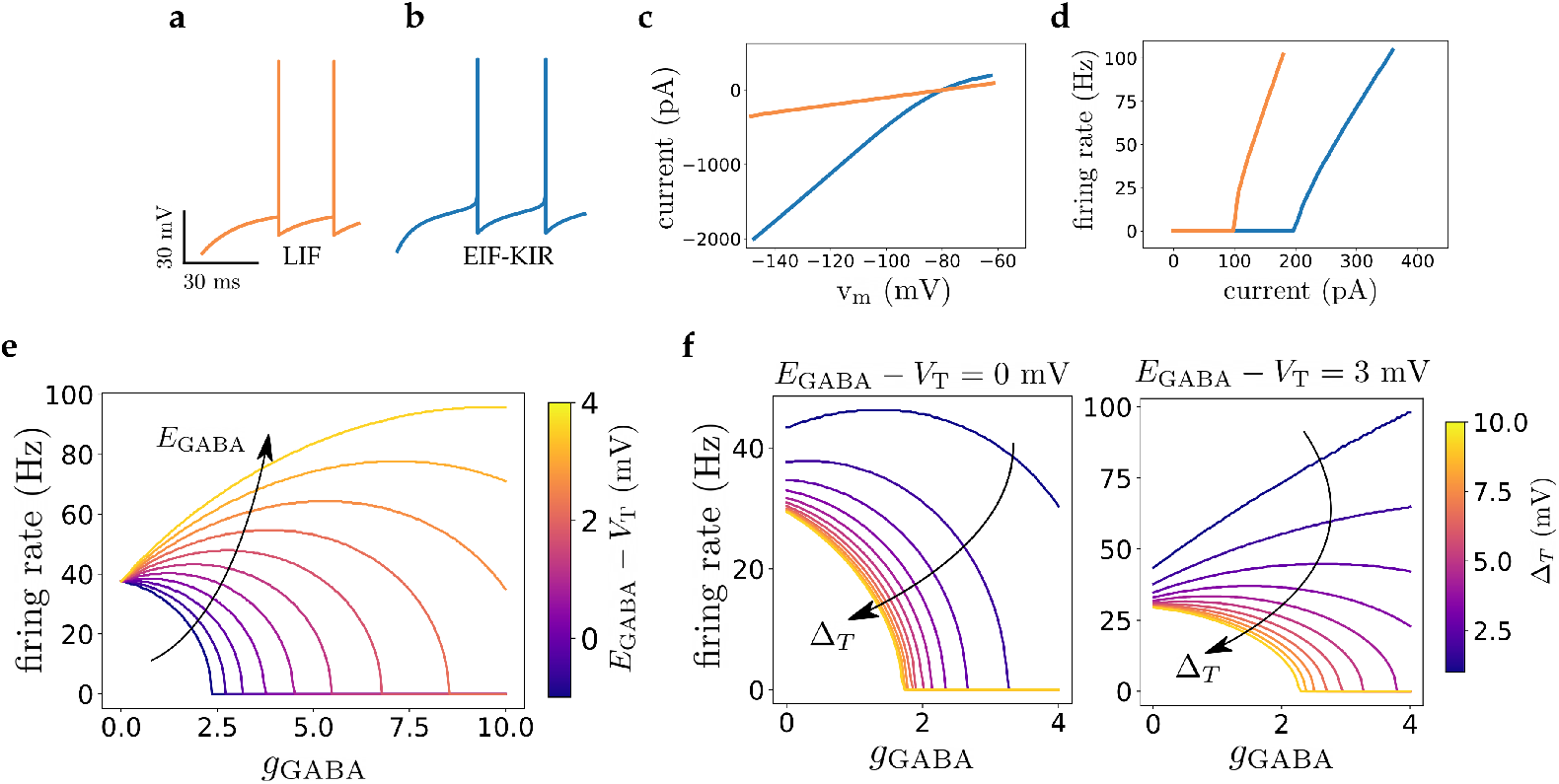
Dynamics of EIF-Kir neuronal model. **a)** Example of membrane potential dynamics of the LIF model of Eq (1) in response to a constant conductance with *g*_Glu_ = 0.4. **b)** Example of membrane potential dynamics of EIF-Kir model of Eq (2) in response to *g*_Glu_ = 0.8. This is similar to LIF model with the addition of a Kir current and a spike generating exponential current. Note the smooth dynamics leading to spike generation in the EIF-Kir model. **c)** V-I curve of the LIF (orange) and EIF-Kir (blue) models. Note the non-linearity of the blue curve due to Kir current. **d)** f-I relationship of the LIF (orange) and EIF-Kir (blue) models. **e)** Firing rate vs GABA conductance in the EIF-Kir model, for different values of GABA reversal potentials and *g*_Glu_ = 0.8. The non-monotonic effect is clearly observed in this model for *E*_GABA_ near *V*_T_, similar to the LIF model. **f)** Effect of the exponential spike generating current on non-monotonicity: Increasing Δ_*T*_ (Eq (2), diminishes non-monotocity for low values of *E*_GABA_ (left) and shifts non-monotonicity for high values of *E*_GABA_ (right), effectively moving non-monotonic regime to higher values of *E*_GABA_.

### Population-level effects of GABA

In the previous sections, we investigated the effect of GABAergic currents at the single neuron level. In this section, we turn to the effects of GABA at the population level. In particular, we ask if similar non-monotonic effects can be reproduced in a recurrent neural network, and what is the distribution of such effects across the population. We choose to focus on a network model of a striatal micro-circuit, for several reasons: striatal SPNs show relatively high GABA reversal potentials [30, 40, 41]; their activity is strongly modulated by the feedforward GABAergic synaptic inputs from FSIs; and several experiments have shown paradoxical effects of FSI feedforward inputs on SPNs [14, 18, 22]. The network consists of three different populations, fast spiking interneurons (FSIs), and direct and indirect spiny projection neurons (dSPNs, iSPNs). There are several other interneuron types in striatum; however, here we only include FSIs as they provide the major GABAergic inputs to SPNs [29, 31].

Fig 6a presents a schematic of the model in which neurons receive uncorrelated glutamatergic Poisson inputs, representing cortical synaptic currents, and are connected by GABAergic synapses. Action potentials of GABAergic neurons generates post-synaptic conductances, *g*_GABA_(*t*), of form Eq 3 shown in Fig 6b. The excitatory inputs are tuned such that the average firing rate of SPNs and FSIs are close to those recorded in awake rodents, which are about 1 Hz and 10 Hz respectively [18]. Fig 6c-d show raster plots and membrane potentials of randomly selected neurons. Membrane potentials fluctuate in the sub-threshold range due to the random arrival of synaptic inputs, leading to irregular firing at low rates. The distribution of membrane potential of SPNs are presented in Fig 6e. It is close to a Gaussian, with a width of 1.1 mV. Fig 6f shows how SPN population mean firing rate depends on FSI population mean rate. When *E*_GABA_ is within a few mV of *E*_thr_, (here 1 mV), a non-monotonic dependence can be observed, while lower *E*_GABA_ results in an average inhibitory effect of FSIs on SPNs, in agreement with the results at the single neuron level. Thus, in these simulations, both increasing or decreasing FSI activity from its normal firing rate (10 Hz) results in inhibition of the SPN population, similar to in vivo observations [18]. The difference between dSPN and iSPN curves are due to asymmetric connectivity of FSIs to these two populations [29]. As FSIs are connected to dSPNs with higher probability, the direct pathway is modulated more strongly by FSI feedforward currents. Thus, these analyses show prominent effects of FSIs on direct-indirect pathway balance. In addition to a change in SPN mean firing rate, FSIs also affect the width of the distribution of SPN firing rates. Fig 6g shows the distributions of SPNs firing rates for two values of FSI activity, 0 Hz and 25 Hz. While the mean rates are similar, the width of the distribution increases with increasing FSI activity. Thus, there is a strong heterogeneity in changes in individual SPNs firing rate, shown in Fig 6h. This heterogeneity can be explained by non-monotonic effects of FSI input on SPNs firing rate. As shown in Fig 6i, SPNs with smaller numbers of FSI inputs tend to increase their firing rate, while SPNs receiving strong FSI inputs tend to decrease their rates. Thus, our network model can explain why manipulations of FSI activity can result in excitatory or inhibitory effect in different subsets of SPNs [14, 22].

**Fig 6.**
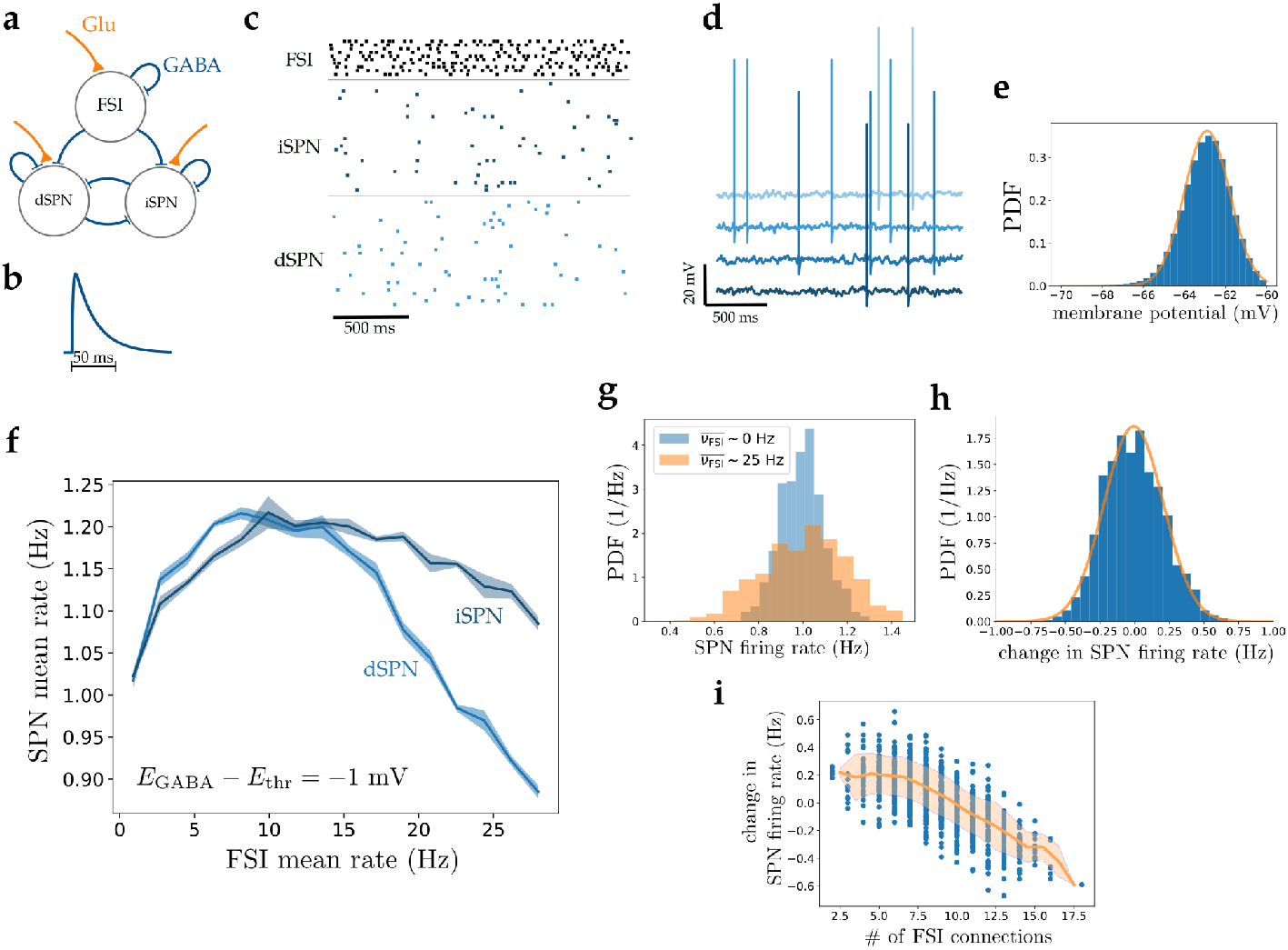
Non-monotonic effects of GABA in a striatal microcircuit model. **a)** Schematic of striatal micro-circuit, composed of three types of neurons, fast spiking interneurons (FSIs) and direct and indirect spiny projection neurons (dSPNs and iSPNs representing direct and indirect pathways). All three cell types receive glutamatergic cortical inputs (orange), and interact through GABAergic synapses (blue). **b)** GABAergic post-synaptic conductance (PSC) triggered by a single pre-synaptic spike (Eq 3). **c)** Raster plot of randomly selected neurons from different populations. External inputs are chosen so that FSIs and SPNs fire at experimentally recorded rates *in vivo* (10 Hz and 1 Hz respectively) **d)** Examples of membrane potentials of randomly selected SPNs. **e)** Probability distribution function of SPNs membrane potential. The orange line is a fitted Gaussian with a standard deviation of 1.1 mV. **f)** Non-monotonic response of SPN population mean firing rate as a function of FSI population mean firing rate, when *E*_GABA_ is 1 mV below *E*_thr_ in SPNs. The shaded area represents standard deviation on the mean, computed from a total of 100 independent network realizations. **g)** Distribution of SPN firing rates when FSI mean firing rate is 0 Hz (blue) and 25 Hz (orange) computed from 100s of network simulations. The mean rates are similar, while the standard deviations of the distributions are 1.1 and 1.9 Hz respectively. **h)** Distribution of the change in firing rate of individual SPNs when FSI rate changes from 0 Hz to 25 Hz. Individual SPNs respond heterogeneously to changes of FSI activity while the mean rate does not change significantly. **i)** Scatter plot of change in SPN firing rate (as described in part g) as a function of number of incoming FSI connections. The change in firing rate of an SPN is strongly correlated with the number of its afferent FSI connections.

## Discussion

GABAergic synaptic currents show hyperpolarizing or depolarizing effects depending on neuronal identity, brain region, and developmental stage. It has also been shown that the characteristics of GABAergic transmission are affected in several pathological conditions. Intuitively, sufficiently low (high) GABA reversal potentials lead to exclusively inhibitory (excitatory) effect of GABA, as GABA drives the membrane potential towards its reversal potential. However, the effect of GABA on neuronal firing rate can be more subtle than this inhibition-excitation dichotomy. In this paper, we used a leaky integrate and fire model to quantify the different effects of GABA and provided a phase diagram of such effects. In particular, we showed that as GABA reversal potential increases, there exists a non-monotonic regime in between purely inhibitory and excitatory regimes, in which small GABAergic currents have an excitatory effect on firing rate, while large ones inhibit the firing rate of the post-synaptic neuron. In other words, in the non-monotonic regime, GABA can be inhibitory or excitatory depending on its input strength. The non-monotonic regime appears when the GABA reversal potential is below, but sufficiently close to the firing threshold. We also studied the effects of GABA in the presence of input noise. We found that the non-monotonic region expands with increasing noise level, showing that fluctuations in synaptic input make the non-monotonic effect stronger and present in a wider range of GABA reversal potentials. We also showed that this non-monotonic effect is qualitatively observed in a more realistic neuronal model that captures some of the electrophysiological properties of striatal spiny projection neurons. Furthermore, using simulations of a network model of local striatal circuit, we showed that non-monotonicity can also be observed at the population level. In addition, we showed that in the network model, the effects of changing FSI firing rates on SPNs are strongly heterogeneous.

To observe non-monotonicity, two conditions should hold: (i) The GABA reversal potential should be sufficiently close to threshold; (ii) The strength of GABAergic inputs should be in a range that encompasses the GABA conductance that maximizes the firing rate (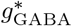 in Fig 2e and Fig 3f). These conditions are potentially satisfied in several neuronal types in physiological conditions. In particular, striatal SPNs [30, 40, 41], fast spiking interneurons in amygdala, cortex [42], and cerebellar interneurons [4] have all been shown to have GABA reversal potentials that are close to the spike threshold.

The non-monotonic effect can be intuitively understood as a competition of two factors. In the deterministic case, the neuronal firing rate depends on GABAergic conductance *g*_GABA_ through two quantities: the effective time constant, *τ_eff_* = *τ*/*g*_eff_, and the effective reversal potential *E*_eff_. On one hand, increasing *g*_GABA_ decreases the effective time constant, leading to potentially faster firing. On the other hand, it decreases the effective reversal potential, leading potentially to a reduction in firing rate. In the non-monotonic regime and when GABAergic conductance is small enough, the effect of *τ*_eff_ shortening wins over that of *E*_eff_ decrease. However, the effect of Eeff wins when the GABAergic conductance is large enough, since in the limit of large *g*_GABA_, the membrane potential converges to *E*_eff_ → *E*_GABA_ < *E*_thr_, and thus never crosses the firing threshold. Another way to understand the effect of GABA in this regime is that when *E*_GABA_ is in between the reset and the threshold, the dynamics of the membrane potential is initially below *E*_GABA_, but then crosses *E*_GABA_ before in turn crossing threshold. Thus, in the initial part of the trajectory GABA tends to speed up the voltage increase towards threshold, while close to threshold it tends to slow it down. The net effect of GABA thus depends on the relative fraction of time spent below and above *E*_GABA_. This argument explains why GABA is initially excitatory when *E*_GABA_ is close enough to threshold, since in that case, GABA is excitatory in most of the trajectory leading to an action potential. However, as the inhibitory conductance increases, Eeff decreases and eventually falls below *E*_thr_, therefore stopping firing altogether.

In the stochastic case, fluctuations in the inputs make it possible for the neuron to fire even when *E*_eff_ is below *E*_thr_. As the amplitude of noise increases, at a parity of output rate, the distribution of membrane potential is pushed towards more hyperpolarized values. Thus, the membrane potential spends an increasing fraction of time below *E*_GABA_ as the GABAergic conductance increases, leading to an enlargement of the non-monotonic regime. Increasing *g*_GABA_ eventually leads to a decrease in firing rate, because of the dampening of the effective noise with increasing conductance (Eq 11).

Our findings at both single neuron and population levels can explain apparently paradoxical effects reported in the experimental literature. In the adult striatum, GABA reversal potentials are believed to be relatively high in spiny projection neurons (SPNs) [30, 40, 41], which could be due to low levels of the KCC2 co-transporter expression in this cell type [43]. This high GABA reversal potential can explain several results in *in vitro* preparations: In particular, why GABA synaptic inputs can sometimes potentiate the response of SPNs to glutamatergic inputs, depending on the relative timing between the two current types [41], and also why pharmacological inhibition of FSIs can reduce such responses [14]. In vivo experiments show even more varied SPN responses to FSI manipulations. One of the most puzzling results come from the study of [18], in which they found that both optogenetic activation and inactivation of FSIs inhibit SPN population activity. While this observed non-monotonicity might be due to experimental artifacts [44] or disynaptic effects [18], our results show that it could also arise due to the non-monotonic dependence of neuronal firing rate on GABA conductance. In particular, this would be consistent with the hypothesis that the striatum is set in a state close to the peak of the firing rate vs GABA conductance curve (see Fig 6f), so that both increasing and decreasing FSI activity leads to a reduction of average SPN firing rates. Note however that other studies have failed to reproduce the reduction of population SPN firing rate when FSI activity is decreased, observed in [18]. In these studies, inhibition of FSIs by pharmacological, chemogenetic and optogenetic methods showed disinhibitory effects on SPN firing in vivo [14, 22]. Importantly, however, these studies also showed that the effects of FSI inhibition on single SPNs are heterogeneous. FSI inhibition leads to both positive and negative SPN activity modulations for different subsets of SPNs - about 60% of SPNs were disinhibited, while the other 40% were inhibited in ref. [22], while this ratio was 74% to 26% in ref. [14]). This strong heterogeneity of effects can be explained in our network model by non-monotonic effects of FSI in which SPNs can be excited or inhibited depending on their FSI GABAergic inputs. Thus, our model reproduces the diversity of effects seen in experimental studies of striatum.

Effects of GABA conductance on neuronal firing rate that differ from pure excitation or pure inhibition have been observed in other cell types and model studies. A non-monotonic dependence of firing rate on GABA conductance has also been found in *vitro* in hippocampal CA1 stratum radiatum interneurons [17]. These authors found that increasing GABAergic conductance first leads to an increase of firing rate, but eventually to a decrease at sufficiently high conductance, and reproduced the effect using a Hodgkin-Huxley type model. Morita et al. used a two-compartmental Hodgkin-Huxley type model, introduced by Wilson [45], with periodic GABAergic synaptic inputs and showed that a similar non-monotonic effect of GABAergic inputs on neuronal firing can be obtained [46, 47]. Wu et al. used a striatal circuit model in which SPNs have an elevated GABA reversal potential, and found that excitatory and inhibitory GABA currents can coexist [48]. In all these models, mixed inhibitory-excitatory effects of GABA are related to a depolarized GABA reversal potential compared to resting membrane potential. Our study complements these previous studies in several ways: First, we used a simplified model that allowed us to analytically characterize the different GABA effects and dissect the mechanisms and parameter space of the non-monotonic effect. Second, we investigated steady states with stationary inputs, rather than transient dynamics or periodic inputs as in some of the previous studies [47]. Third, we systematically analyzed the role of input noise and how it influences the effects of GABA on neuronal firing. Additionally, we speculate that as many neuronal types in several brain regions undergo a reduction in GABA reversal potential through development and neurogenesis [1, 8], this non-monotonic effect should have important effects at specific developmental stages and may play a role in network formation and neural integration into an existing network. Indeed, an excitatory-inhibitory dual role of GABA has been experimentally observed in immature rat hippocampus and neocortex [49, 50].

While the simplicity of our models provide insight into the mechanisms underlying paradoxical GABA effects, our models also have a number of limitations. Parameters in the single neuron models that are considered constants, such as *E*_thr_ or *E*_GABA_, are rather dynamic and depend on complex conductance and ion concentration dynamics. In this paper, we considered steady conductances and neglected the spatiotemporal complexity of GABAergic and glutamatergic currents. GABAergic synaptic currents lead to changes in Cl^−^ concentration due to passage of chloride ions through GABA receptor-channels, and consequently changes in the effective GABA reversal potential [51, 52]. It has also been been shown that in spatially extended models, GABAergic currents can have complex interactions with other inputs depending on their respective locations, which can alter their effects [53, 54]. While our model does not directly address these complex interactions, we believe it provides a good first-order approximation when the time scale of these effects is much longer than the membrane time constant, *τ*. For instance, Cl^−^ concentration in SNr neurons changes on time scales of ~ 1s [54]. In these cases, GABA reversal potential can be considered effectively constant on faster time scales. Our model predicts that small changes of *E*_GABA_, when close to *E*_thr_, can shift GABAergic currents inhibitory-excitatory role. These small shifts could be physiologically obtained by different ionic mechanisms such as activity-dependent Cl^−^ accumulation [52–58] and chloride transporter alterations [59, 60]. Another simplification is that in the population model, we use a simplified binary distribution of synaptic weights (either zero or *G_ij_* in Eq 4). In reality, synaptic weights exhibit broad distributions. A wider distribution of weights would result in higher temporal fluctuations of synaptic inputs, which would then enlarge the non-monotonic region. Heterogeneity of synaptic weights also leads to a broader distribution of synaptic conductances from neuron to neuron, potentially increasing the heterogeneity of GABA effects that we observed in our network model.

Previous works have also explored the potential functional consequences of high GABA reversal potentials. Shunting inhibition has been shown to affect the gain of the input/output neuronal transfer function [61, 62]. GABAergic reversal potentials also have a strong effect on synchronization properties of GABAergic neurons, as shown by several studies [20, 21]. In particular, Vida et al. [20] showed that GABA reversal potentials that are higher than resting membrane potentials lead to oscillation generation with smaller excitatory drive, compared to hyperpolarizing inhibition. In addition, they showed that network oscillations in such networks are more robust to heterogeneity in excitatory drive. During development, depolarizing GABAergic currents have been implicated in the generation of rhythmic activity in neonatal hippocampus [63], and in thalamic reticular nucleus [19]. Studies on epileptic patients point to excitatory GABA effects in several brain regions that contribute to this pathological condition [64–68]. Our results suggest potential additional mechanisms contributing to these effects by providing detailed analysis of GABAergic inputs and their relevant variables in several neuronal models.

In conclusion, the results presented in this paper can explain several experimentally observed paradoxical effects of GABAergic synaptic currents and provide a framework to classify the different effects of GABA on neuronal firing rate, as a function of single neuron and network parameters. We suggest that the dichotomous framework of inhibition-excitation for GABAergic currents does not capture the full spectrum of GABA effects, as it ignores non-monotonic effects that could potentially be relevant in several brain regions. These findings have potential implications for understanding brain development, neural network formation and neural dynamics, in both normal and pathological conditions.

## Acknowledgments

This work was supported by NIH BRAIN R01NS110059.

